# Metabolic Reprogramming by Histone Deacetylase Inhibition Selectively Targets NRF2-activated tumors

**DOI:** 10.1101/2023.04.24.538118

**Authors:** Dimitris Karagiannis, Warren Wu, Albert Li, Makiko Hayashi, Xiao Chen, Michaela Yip, Vaibhav Mangipudy, Xinjing Xu, Francisco J. Sánchez-Rivera, Yadira M. Soto-Feliciano, Jiangbin Ye, Thales Papagiannakopoulos, Chao Lu

## Abstract

Interplay between metabolism and chromatin signaling have been implicated in cancer initiation and progression. However, whether and how metabolic reprogramming in tumors generates specific epigenetic vulnerabilities remain unclear. Lung adenocarcinoma (LUAD) tumors frequently harbor mutations that cause aberrant activation of the NRF2 antioxidant pathway and drive aggressive and chemo-resistant disease. We performed a chromatin-focused CRISPR screen and report that NRF2 activation sensitized LUAD cells to genetic and chemical inhibition of class I histone deacetylases (HDAC). This association was consistently observed across cultured cells, syngeneic mouse models and patient-derived xenografts. HDAC inhibition causes widespread increases in histone H4 acetylation (H4ac) at intergenic regions, but also drives re-targeting of H4ac reader protein BRD4 away from promoters with high H4ac levels and transcriptional downregulation of corresponding genes. Integrative epigenomic, transcriptomic and metabolomic analysis demonstrates that these chromatin changes are associated with reduced flux into amino acid metabolism and *de novo* nucleotide synthesis pathways that are preferentially required for the survival of NRF2-active cancer cells. Together, our findings suggest that metabolic alterations such as NRF2 activation could serve as biomarkers for effective repurposing of HDAC inhibitors to treat solid tumors.

## Introduction

Eukaryotic cells have evolved sophisticated mechanisms to sense and integrate extracellular information into an intrinsic signaling system that regulates transcription so that environmental fluctuations can be delivered and responded to in a timely and accurate manner. A key player in the process is chromatin, as many chromatin-modifying reactions require not only the proteinaceous enzymes, but also small-molecule substrates/co-factors that are intermediates of central carbon metabolism. Indeed, it has been well-documented that chemical modifications of DNA and histones can act as sensors for fluctuations in cellular metabolic flux and in turn mediate the transcriptional response to maintain metabolic homeostasis^1^. Importantly, we and others have reported that these mechanisms can be hijacked by cancer cells to reprogram gene expression and facilitate tumor progression^2–4^. It is less clear, however, if distinct metabolic abnormalities also render cancer cells vulnerable to perturbations of chromatin regulatory mechanisms. As a result, the therapeutic potential of targeting the crosstalk between chromatin and metabolism remains under-explored.

Nearly 20% of lung adenocarcinoma (LUAD) tumors carry loss-of-function mutations in *KEAP1* or gain-of-function mutations in *NFE2L2* genes, both of which lead to activation of the NRF2 antioxidant response pathway^5^. Activation of this pathway conveys several tumor-promoting properties to cells, including an increase in anabolic processes, production of antioxidants and detoxifying enzymes^5–7^. These effects promote aggressive disease and drug resistance, which makes NRF2-active tumors particularly hard to treat. Aberrant NRF2 activation also occurs in other cancer contexts, such as head and neck squamous cell carcinoma and hepatocellular carcinoma, through genetic and non-genetic mechanisms^8, 9^. It has been reported that metabolic reprogramming upon NRF2 activation confers sensitivity to glutaminase inhibition^10, 11^. However, the KEAPSAKE clinical trial (NCT04265534), which evaluated the efficacy of glutaminase inhibitor CB-839 in patients with *KEAP1* mutation, was discontinued due to lack of clinical benefit. As a result, identifying selective vulnerabilities of NRF2-active cancers that can be exploited for more effective treatment remains a key challenge.

Similar to many cancer-associated metabolic alterations, previous reports have linked NRF2 activation to dysregulated chromatin state^12, 13^. However, little is known about the underlying mechanisms and their importance for therapy. In this study, we sought to test if metabolic reprogramming by NRF2 activation confers potential chromatin-based vulnerabilities. Through a chromatin-focused CRISPR-Cas9 genetic screen, we uncovered an NRF2-driven sensitivity to Class I histone deacetylase (HDAC) inhibition and defined the underlying molecular basis using integrative epigenomic, transcriptomic and metabolomics analysis. Our findings suggest that cancer cells harboring metabolic alterations may exhibit strong and specific dependencies on chromatin regulators that can be therapeutically exploited and highlight the potential of combinatorial targeting of metabolism and chromatin – two emerging and intimately linked cancer molecular hallmarks.

## Results

### NRF2 activation confers preferential vulnerability to loss of Class I HDACs

To model and study NRF2 activation in LUAD, we used mouse lung adenocarcinoma cell lines derived from tumors generated through a genetically engineered mouse model (GEMM) of *Kras^G12D/+^*;*p53^-/-^*driven lung adenocarcinoma. This system also enables CRISPR-Cas9 mediated knockout (KO) of a gene of interest such as *Keap1*^14^. As previously described, *Kras^G12D/+^;p53^-/-^* (KP) and *Kras^G12D/+^*;*p53^-/-^*;*Keap1^-/-^* (KPK) tumors were generated using sgRNAs against *tdTomato* (non-targeting control) or *Keap1*, respectively^10^. Thus, KPK cell lines represent tumor-derived cells with constitutively activated NRF2 pathway and KP cell lines serve as control cells with normal NRF2 activity.

To identify novel chromatin vulnerabilities associated with NRF2 activation, we performed a targeted CRISPR-Cas9 genetic screen. KP and KPK cells were infected with a pool of single-guide RNAs (sgRNAs) targeting 612 chromatin regulators^15^ and passaged for 14 population doublings (**Figure 1A**) . To confirm the quality of the screen, we compared the gene effect scores determined in this screen with that from genome-wide CRISPR-Cas9 screens in non-small cell lung cancer (NSCLC) cell lines from the DepMap database^16, 17^ and found strong correlation between the two datasets (**Figure 1B**). Genes more essential in the context of activated NRF2 were identified by comparing the decreases in the abundance of sgRNAs in KP vs. KPK cells over time (**Supplementary Table 1**). Notably, among genes that were essential in KP but not KPK cells the top hit was *Ube2m* (**Figure S1A**), which is known to interact with the KEAP1 and Cullin-RING ligase (CRL) E3 ligase complex, and thus served as a positive control in our screen. Analysis of significantly depleted genes revealed multiple differential dependencies, including genes encoding Class I HDACs *Hdac1*, *Hdac2* and *Hdac3* which were synthetic lethal with *Keap1* loss (**Figures 1C-D, S1B**). Consistent with this result, in a competition assay assessing the fitness of cell populations carrying various sgRNAs targeting HDAC genes, KPK cells were more sensitive than KP cells to the loss of HDAC1-3 **(Figures 1E-G)**. Taken together, our results suggest that Class I HDAC genes are preferentially required for KPK cell viability and represent candidate therapeutic targets in the context of NRF2 activation.

**Figure 1.**
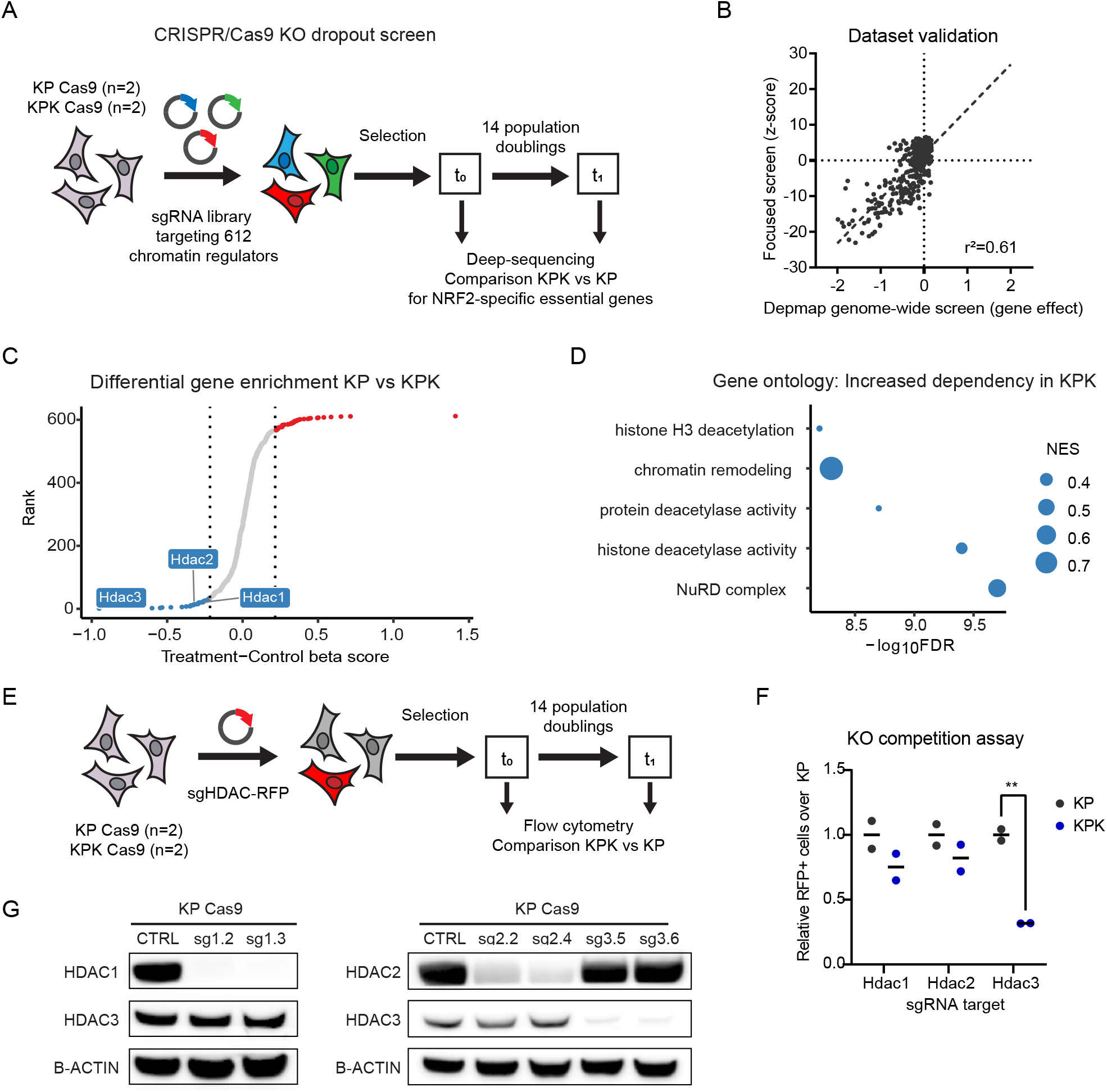
NRF2 activation confers preferential vulnerability to loss of Class I HDACs. **A.** Schematic of chromatin focused CRISPR/Cas9 genetic screen. **B.** Pearson correlation of gene z-score (KP cells t1 vs t0) of focused CRISPR/Cas9 screen to gene effect score of NSCLC cell lines (n=94) from CERES 21Q3 DepMap dataset. **C.** Scatter plot of genes ranked by beta-score (KPK-KP) and ontology analysis **(D)** of genes more essential in KPK cells. **E**. Schematic of competition assay employed to validate hits from the CRIPSR screen. **F**. Competition assay in KP (n=2) and KPK (n=2) cells to CRISPR/Cas9-mediated KO of HDAC genes and HDAC protein levels upon KO (**G**).

### NRF2 activation confers HDAC inhibitor sensitivity

To further validate the genetic screen results and investigate the therapeutic potential, we examined the effect of pharmacologic inhibition of Class I HDACs on the viability of cells with activated NRF2. We also used another isogenic system with the overexpression of NRF2ΔNeh2, a gain-of-function truncated NRF2 mutant lacking the KEAP1 interacting domain^18^ **(Figure 2A)**. NRF2ΔNeh2 robustly induced NRF2 activation in KP cells, as indicated by stabilization of NRF2 protein levels and induction of NRF2 target gene expression (**Figures 2B-C**). Consistent with the association between NRF2 activation and genetic dependency on Class I HDACs, NRF2ΔNeh2-expressing KP cells were more sensitive to the FDA-approved Class I HDAC inhibitor Romidepsin^19^ *in vitro,* relative to their KP controls (**Figure 2D)**. To ensure that this finding is not due to selective pressures of long-term NRF2 activation, we employed three additional experimental systems. To induce transient NRF2 activation, we either used KI696, a small molecule that disrupts the interaction between KEAP1 and NRF2, or a dox-inducible system of NRF2ΔNeh2 overexpression. In both cases, we observed increased sensitivity to Romidepsin treatment. Moreover, we overexpressed KEAP1 in KPK cells and observed reduced NRF2 protein levels and resistance to Romidepsin **(Figures 2E-F**). Additionally, Romidepsin significantly suppressed the *in vivo* growth of KP tumors overexpressing NRF2ΔNeh2, but not control tumors (**Figures 2G-H**).

**Figure 2.**
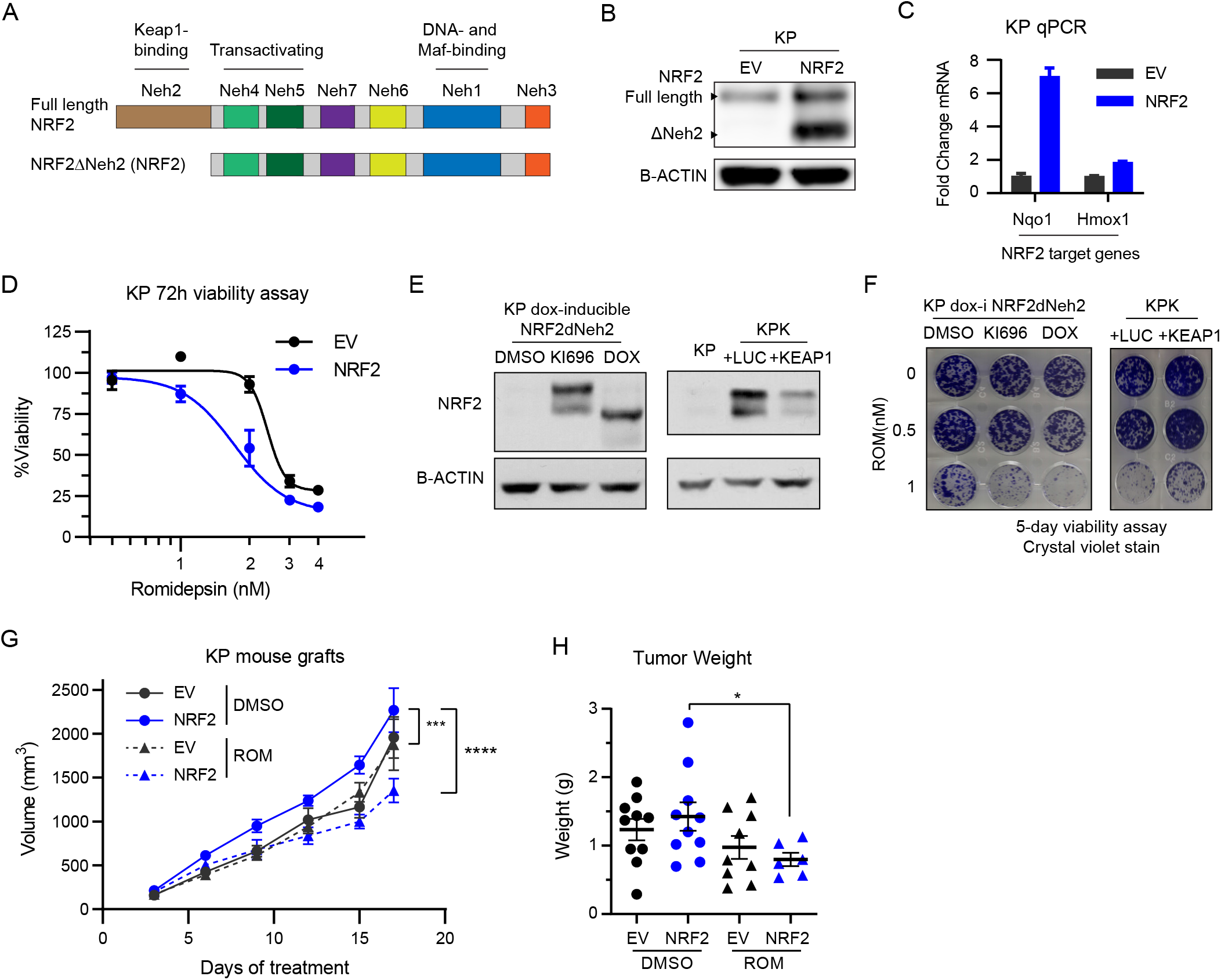
NRF2 activation confers HDAC inhibitor sensitivity. **A.** Schematic indicating domains of the full-length and the ΔNeh2 over-active mutant isoforms of NRF2 protein¹. **B.** NRF2 protein levels upon of overexpression of NRF2ΔNeh2 in KP cells (NRF2). **C.** Expression of NRF2 target genes upon overexpression of NRF2ΔNeh2 in KP cells. **D.** Viability assay of KP cells treated with the indicated concentrations of Romidepsin for 72h. **E.** NRF2 protein levels of KP cells carrying dox-inducible NRF2ΔNeh2 upon treatment with DMSO, KI696 or doxycycline (DOX) and of KPK cells overexpressing luciferase (LUC) or KEAP1. **F.** Viability assay of KP and KPK cells treated with the indicated concentrations for 5 days. **G.** Growth of subcutaneous KP tumors in C57/BL6 mice treated with Romidepsin or vehicle, and tumor weight at the end of treatment (**H**).

To test if this finding can be extended to other HDAC inhibitors, we used four additional HDAC inhibitors: the Class I HDAC inhibitors Entinostat and ACY-957, and the pan-HDAC inhibitors Vorinostat and Belinostat. Both in our cell lines, as well as in NSCLC cell lines from the DepMap dataset, NRF2 activation conferred sensitivity to Class I-specific inhibitors but not pan-HDAC inhibitors (**Figures S2A-B)**. HDAC inhibitors are associated with DNA damage and programmed cell death^20–22^. To assess apoptosis and DNA damage levels, we looked at phosphorylation of H2A.X (γH2AX) and cleaved caspase 3 upon Romidepsin treatment at a concentration where NRF2-activated cells showed increased sensitivity (**Figure S2C**). Results indicate modest increase in DNA damage and apoptosis, which were similar between EV and NRF2 cells, suggesting that the observed differences in Romidepsin sensitivity are not due to increased DNA damage and/or apoptosis (**Figure S2D**).

### Romidepsin alters gene expression by genomic redistribution of histone acetylation and BRD4

Class I HDACs are major regulators of histone acetylation. To investigate how HDAC inhibition affects histone acetylation to reprogram gene expression, we performed CUT&Tag^23^ upon Romidepsin treatment to profile genomic distribution of histone acetylation marks H3K27ac and H4ac (poly-acetylation on H4K5, H4K8, H4K12, H4K16), as well as BRD4, a histone acetylation reader protein that activates gene transcription, in duplicate (**Figure S3A**). We also performed RNA-sequencing and integrated the epigenomic datasets to correlate changes in histone acetylation landscape to differential gene expression.

HDAC inhibition with Romidepsin induced broad gene expression changes including decreased expression of 1,692 genes (**Figure 3A**), which were highly concordant between control and NRF2-active cells (**Figure S3B**). As expected, Romidepsin induced global increase in histone acetylation levels, while BRD4 levels were largely unchanged (**Figure S3C**). We first assessed how genome-wide histone acetylation and BRD4 distributions were affected by HDAC inhibition. We annotated genomic features of H4ac and BRD4 peaks and found that Romidepsin induced a redistribution of peaks from promoters to distal intergenic regions (**Figure 3B**). Consistently, H4ac signal and BRD4 binding at promoter-associated peaks were reduced following Romidepsin treatment (**Figure 3C**). Furthermore, we measured H4ac reads at peaks vs. random genomic regions and observed that upon Romidepsin treatment H4ac was reduced at peaks and increased in random regions (**Figure 3D**). Indeed, Romidepsin treatment led to a >2-fold decrease in the ratio of H4ac peak signal over genome average (**Figure 3D**). Finally, we found that the initial levels of H4ac and BRD4 enrichment correlated with the degree of loss in H4ac and BRD4 binding following Romidepsin treatment (**Figures 3E, S3D-E**). Together, these results suggest a model where HDAC inhibition alters the ratio between the abundance of H4ac at promoter-associated peaks and at distal intergenic regions, which in turn dilutes BRD4 binding at H4ac-high promoters (**Figure S3F**).

**Figure 3.**
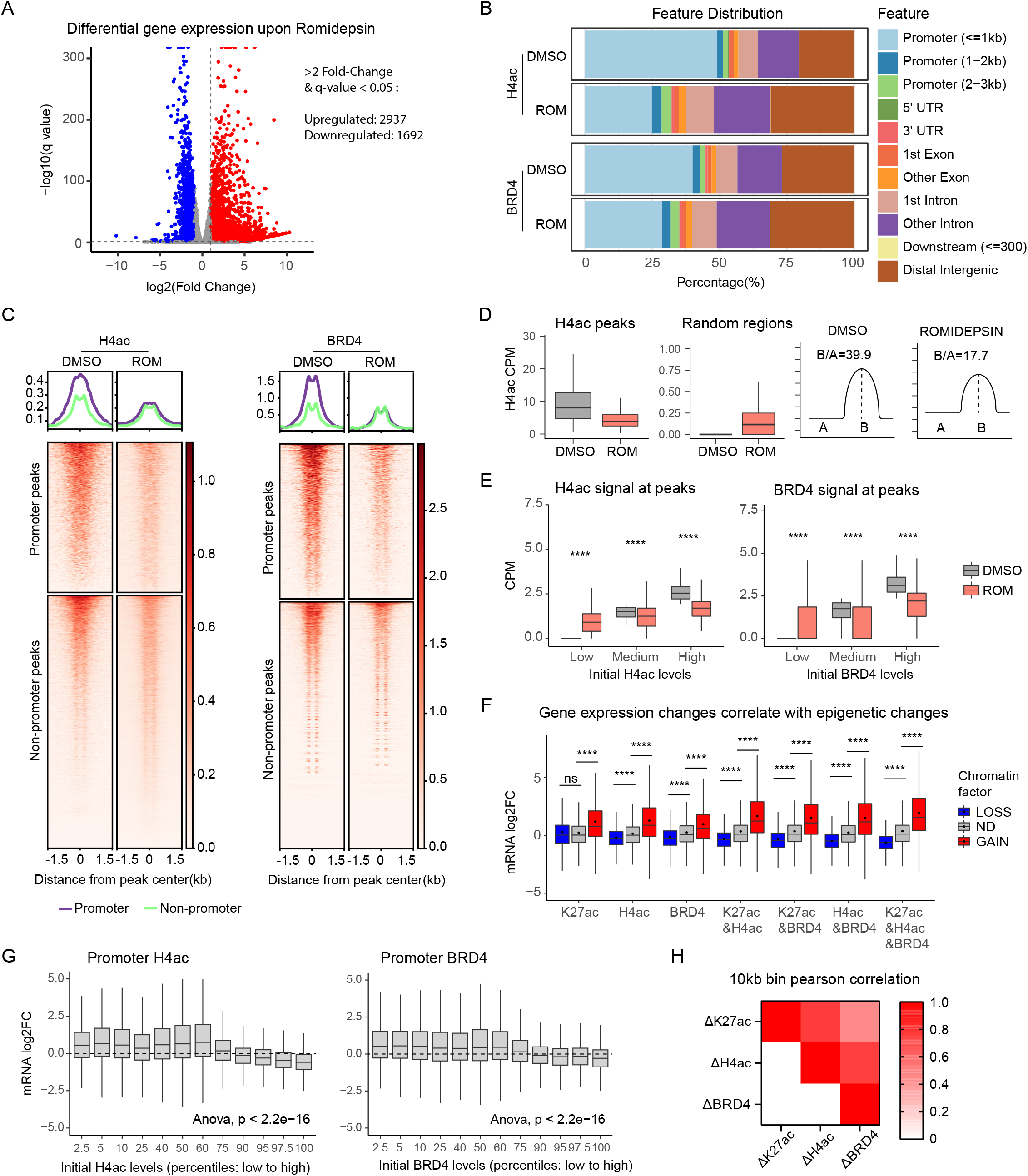
Romidepsin alters gene expression by genomic redistribution of histone acetylation and BRD4. **A.** Volcano plot of differential gene expression upon Romidepsin treatment (5nM, 16h) of KP NRF2 cells. **B**. Distribution of called H4ac and BRD4 peaks (n=2) across genomic features. **C**. Heatmaps of histone acetylation signal at peaks that do or do not overlap with promoters (within 2.5 from a TSS). **D**. Box plots of read CPM at called peaks or random regions and illustrations indicating the ratio of signal at peaks compared to background (random regions). **E.** Box plots of read counts per million (CPM) at consensus peaks with low, medium or high levels at DMSO treatment (cutoff: peaks were ranked by CPM and split into thirds; one out of two replicates shown). **F**. Fold-change expression (ROM/DMSO; DEseq2 analysis of RNA-seq) of genes that gain, lose, or have no difference (ND) in promoter histone acetylation and/or BRD4 binding (cutoff: 1.5-fold change). **G**. Fold-change expression (as in **F**) of genes with variable initial levels of H4ac and BRD4 binding (promoters were ranked by CPM and split into percentiles; average of two replicates shown). **H**. Pearson correlation of genome-wide signal difference (ROM-DMSO) split into 10kb bins between histone acetylation marks and BRD4 (mean of two replicates).

Importantly, these relative changes in histone acetylation, particularly H4ac, and BRD4 binding at gene promoters correlated strongly with changes in gene expression (**Figure 3F**). Consistently, genes with high levels of H4ac and BRD4 binding at promoters also showed the largest degree of decrease in expression following HDAC inhibition (**Figure 3G**). We also measured absolute (with spike-in normalization) changes in H3K27ac and H4ac and found that they poorly correlated with gene expression changes: when adjusting histone acetylation signal based on total abundance, there was increased histone acetylation associated with both upregulated and downregulated genes (**Figure S3G**). Therefore, the relative changes in H4ac appear to be a major driver of BRD4 re-targeting and transcriptomic changes.

Overall, these results suggest that Romidepsin induces gene expression changes, primarily as a result of broad redistribution of H4ac and BRD4 binding. In particular, we find diffusion of H4 acetylation away from promoters and highly acetylated peaks. Previous report suggests that BRD4 distribution is affected similarly to relative H4ac changes, including displacement from gene promoters^24^. Indeed, we found that genome-wide changes in BRD4 binding correlated well with H4ac changes and to a less degree H3K27ac (**Figure 3H**). This suggests a model where HDAC inhibition induces redistribution of BRD4 driven by changes in relative H4ac levels and alters gene expression (**Figure S3F**).

### Romidepsin regulates expression of genes that represent known and novel metabolic vulnerabilities of NRF2-active cells

Since Romidepsin induced similar transcriptomic changes in control and NRF2 active cells (**Figure S3B**) but showed preferential toxicity towards NRF2 active cells, we reasoned that the differentially expressed genes could affect pathways that are more essential for NRF2-active cell viability. To explore this hypothesis, we first examined if known vulnerabilities of KEAP1 loss and NRF2 activation are transcriptionally regulated by Romidepsin. It has been well documented that NRF2 activation is associated with a specific dependency on glutamine uptake and catabolism^10, 11, 25^. In addition, NRF2 activation has been shown to promote serine and glycine biosynthesis and dependency^26, 27^. Therefore, we examined expression of genes involved in glutamine uptake^28^/metabolism and serine/glycine biosynthesis pathway. We found that Romidepsin induced downregulation of these genes *in vitro* (**Figures 4A**, **S4A**). Moreover, Romidepsin treatment *in vivo* led to a reduction in protein levels of ATF4, a master transcriptional regulator of amino acid metabolism (**Figures S4B-C**). In contrast, we did not find consistent gene expression changes in glycolysis, TCA cycle and NRF2 target genes (**Figure 4A**).

**Figure 4.**
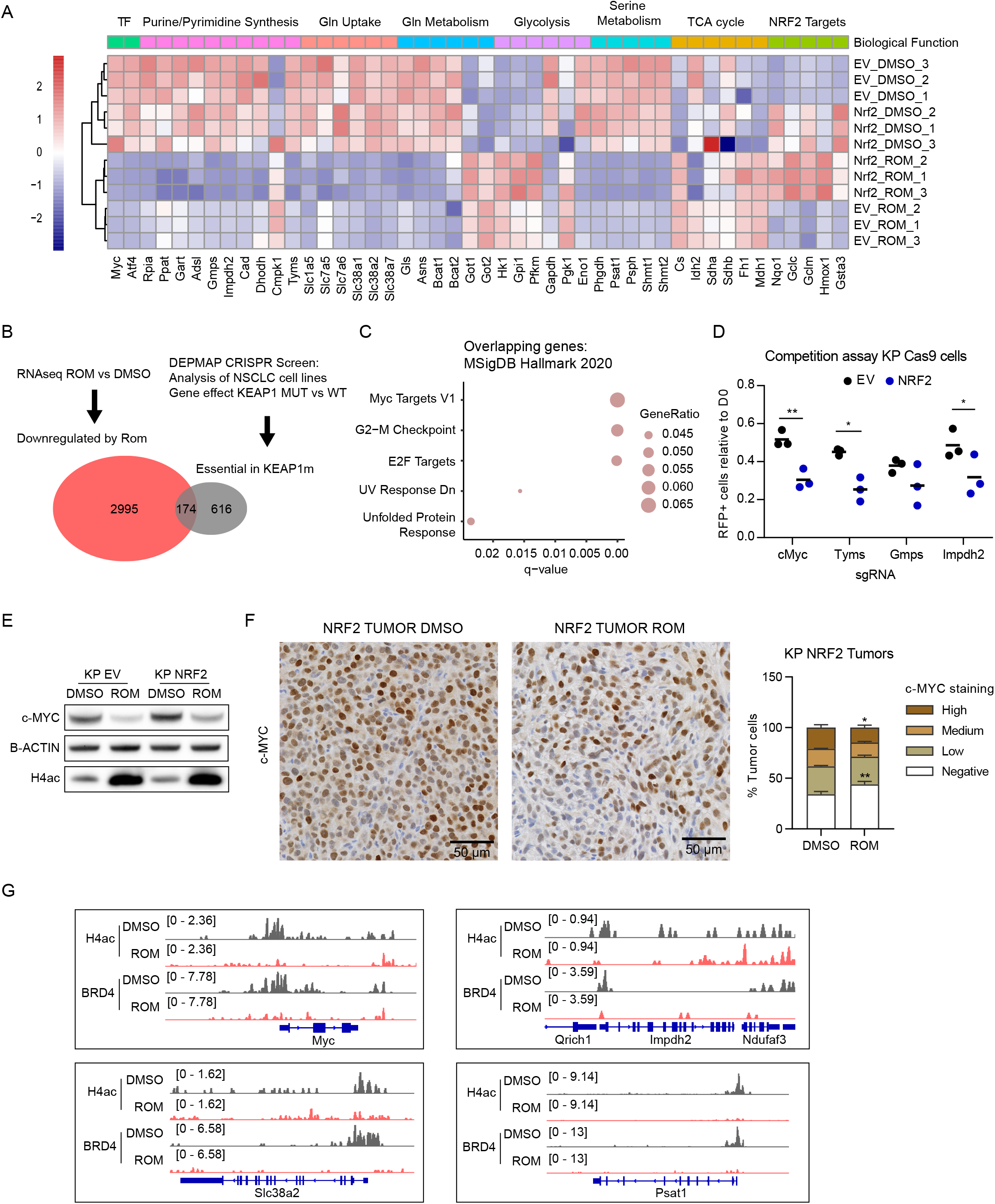
Romidepsin regulates expression of genes that represent known and novel metabolic vulnerabilities of NRF2-active cells. **A**. Heatmap showing expression of metabolic genes (RNA-seq; normalized by DEseq2 and scaled for each gene). **B**. Venn diagram showing overlap between genes downregulated by Romidepsin (cutoff: 1.5-fold change and FDR<0.05) in KP NRF2 cells and genes that are more essential in KEAP-mutant NSCLC cell lines from the CERES DepMap dataset (cutoff: 0.05 gene effect difference), and MsigDB Hallmark ontology of the overlapping genes (**C**). **D**. Competition assay showing fraction of RFP+ KP Cas9 cells after 14 population doublings, indicating percentage of cells that carry sgRNA against the indicated genes (n=3). **E**. Western blot showing levels of c-MYC and H4ac upon Romidepsin treatment of KP cells (1nM for 24h). **F**. Immunohistochemistry (IHC) staining of c-MYC and quantitation in KP NRF2 tumors after 17 days of treatment with DMSO or Romidepsin (related to figure 2G-H) **G**. Bedgraphs of H4ac and BRD4 binding at indicated gene loci.

We next analyzed the CERES genome-wide CRISPR screen dataset of human cell lines from the DepMap database^16, 17^. We identified genes that represent specific dependencies for *KEAP1*-mutant NSCLC cell lines and intersected them with the genes downregulated by Romidepsin (**Figure 4B**). Gene ontology revealed enrichment in MYC targets (**Figure 4C**), including *Myc* itself and its target genes involved in purine and pyrimidine synthesis (**Figure S4D**). Using a competitive cell proliferation assay, we confirmed that genetic knock-out of *Myc* and several *de novo* nucleotide synthesis genes was more detrimental to the survival of KP cells with NRF2 activation (**Figurse 4D, S4E**). Furthermore, Romidepsin reduced MYC protein levels *in vitro* and *in vivo* **(Figures 4A, 4E-F, S4F**).

Taken together, our results suggest that Romidepsin induces downregulation of several metabolic genes that are more essential for the survival of cells with NRF2 activation. Importantly, the majority of these transcriptional changes correlated with direct loss of promoter H4ac and BRD4 binding (**Figures 4G, S4G**), while the rest are likely a downstream effect of MYC suppression. Therefore, these findings suggest that Romidepsin induces epigenetic and transcriptional changes that target known and novel NRF2-specific metabolic vulnerabilities.

### Romidepsin disrupts metabolic processes that are essential for NRF2-active cells

To examine how Romidepsin-induced changes in metabolic gene expression affect metabolic flux, we performed targeted metabolite tracing analysis. KP cells with or without NRF2 activation were treated with DMSO, Romidepsin or the glutaminase inhibitor CB-839 for 24 hours, and then cultured in ^13^C-glucose for 1 and 24 hours, or ^13^C-glutamine for 8 hours before harvesting (**Figures 5A-B, S5A**). Glutamine tracing indicated that in Romidepsin-treated but not CB-839- treated cells, the proportion of ^13^C-labeled glutamine was reduced, indicating reduced glutamine uptake (**Figure 5C**). Consistent with this finding, media glutamine consumption was reduced upon Romidepsin treatment (**Figure S5B**). Romidepsin treatment also reduced ^13^C incorporation into further steps of glutamine metabolism (**Figure 5D**; M+5 glutamate, M+3 αKG). Interestingly, in NRF2-active cells, Romidepsin and CB-839 had a comparable effect on reducing glutamine-derived ^13^C incorporation into TCA cycle metabolites (**Figure 5E;** M+4 citrate, M+4 succinate, M+4 fumarate, M+4 malate). In addition, we found strongly reduced incorporation of glutamine in pyrimidine nucleotides (**Figure 5F**; M+1, +2 and +3 UTP), which suggests a disruption of *de novo* nucleotide synthesis.

**Figure 5.**
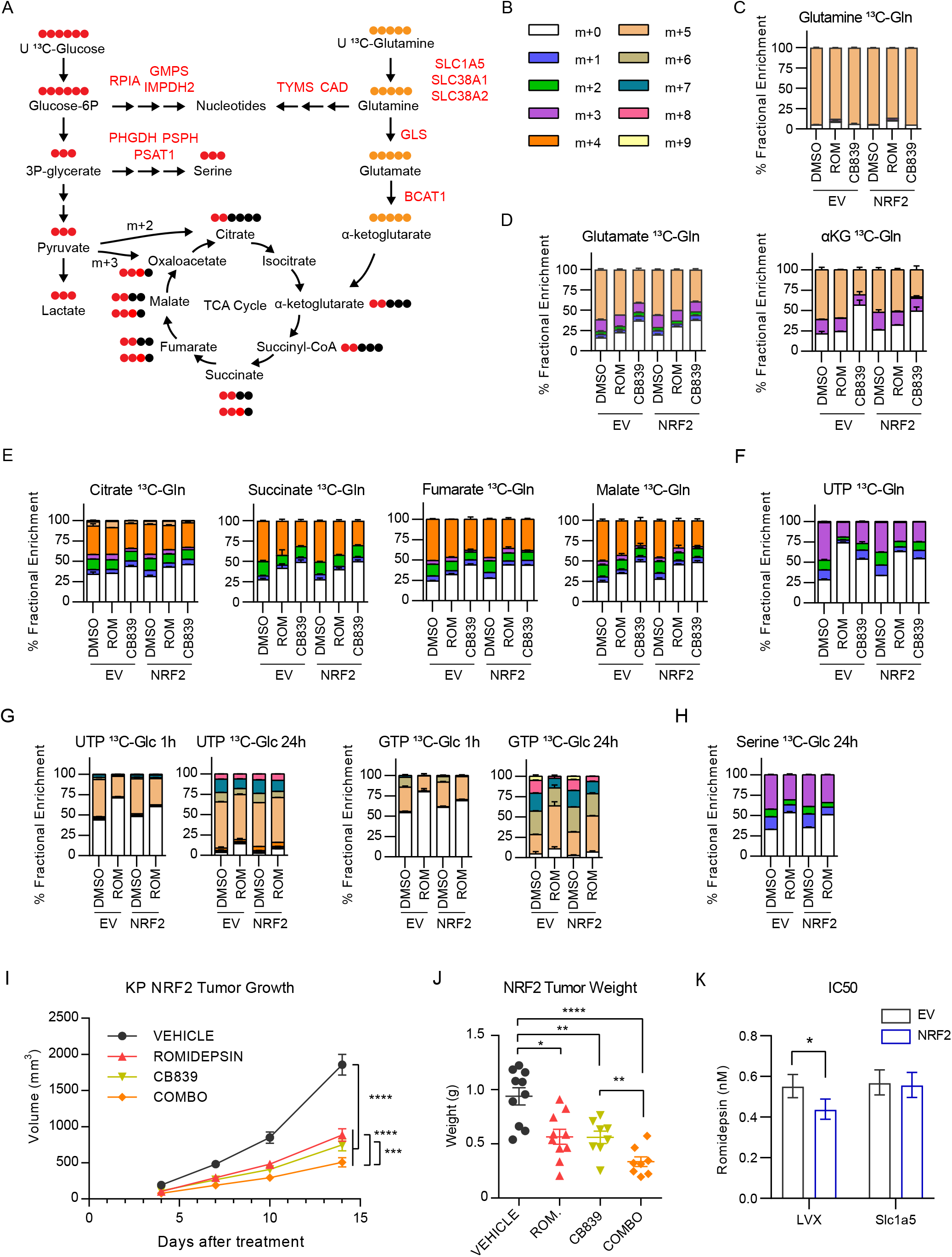
Romidepsin disrupts metabolic processes that are vulnerable in NRF2-active cells. **A.** Schematic indicating glucose and glutamine metabolic processes, labeled carbon contribution and genes that are downregulated upon Romidepsin treatment. **B**. Color indications of isotopologues. **C-F**. Bar graphs showing isotopologue percentages for the indicated metabolites after U-¹³C glutamine tracing for 8h. **G-H**. Bar graphs showing isotopologue percentages for the indicated metabolites after U-¹³C glucose tracing for the indicated time. **I**. Growth of subcutaneous KP NRF2 tumors in C57/BL6 mice treated as indicated, and tumor weight at the end of treatment (**J**). Romidepsin IC50 of KP cells transduced with empty vector (LVX) or overexpressing Slc1a5 (SLC1A5); error bars represent 95% confidence interval.

While glycolysis has been reported to be suppressed by HDAC inhibition in other tumor models^29, 30^, we didn’t find this to be the case in our experimental system (**Figure S5D-E**), in agreement with our gene expression data (**Figure 4A**). However, we did find reduced incorporation of glucose-derived ^13^C in purine and pyrimidine nucleotides (**Figure 5G**; 1h M+5 UTP, 24h M+6 and +7 UTP, 1h M+5 and +6 GTP, 24h M+7, +8 and +9 GTP), again consistent with the gene expression results, and indicates reduced rate of *de novo* nucleotide synthesis. Moreover, we found reduced incorporation of glucose-derived ^13^C into serine, which suggests a possible defect in serine biosynthesis (**Figure 5H**; M+1, +2 and +3).

Overall, these metabolomic results suggest that Romidepsin disrupts key metabolic processes (serine/glutamine metabolism and *de novo* nucleotide synthesis) that support viability in NRF2- active cells. Specifically, Romidepsin disrupts glutamine flux into the TCA cycle in a similar manner to glutaminase inhibition, which underlies the NRF2-specific sensitivity to glutaminase inhibitor^11^. In agreement with this notion, Romidepsin treatment *in vivo* showed similar efficacy to CB-839 (**Figures 5I-J**). Interestingly, the combination of Romidepsin and CB-839 resulted in improved tumor response than either treatment alone (**Figures 5I-J**), without a significant effect on mouse weight (**Figure S5G**). As another functional validation, we found that overexpression of Slc1a5, a transporter that mediates glutamine uptake, partially rescued sensitivity of NRF2 activated cells to Romidepsin treatment (**Figures 5K, S5H-I**), in support of the hypothesis that impaired glutamine uptake contributes to the NRF2-specific sensitivity to HDAC inhibition.

### NRF2-activation confers sensitivity to HDAC inhibition in human cells and patient-derived xenografts (PDX)

To test if the association between NRF2 activation and HDAC inhibitor sensitivity is conserved in human cells, we used the *KEAP1*-mutant LUAD cell line A549 and targeted *NFE2L2* using the CRISPR/Cas9 system, which led to a partial reduction in NRF2 protein levels and NRF2 target gene expression (**Figures 6A, B**), without affecting proliferation *in vitro*. Consistent with our findings in the murine cell lines, Romidepsin suppressed expression of *MYC*, *ATF4* and several genes involved in *de novo* nucleotide synthesis, glutamine transport and serine synthesis (**Figures 6C, D**). Moreover, NRF2 WT A549 cells were more sensitive to Romidepsin than NRF2 KO cells *in vitro* (**Figure 6E**) and *in vivo* (**Figure 6F)**. Additionally, induction of NRF2 activation in the *KEAP1* wild-type cell line NCI-H2009 cell line by overexpression of KEAP1^R470C^, a dominant negative mutant form of KEAP1^31^, led to increased sensitivity to Romidepsin (**Figure S6A, B**). Finally, in patient-derived xenografts of *KEAP1* wild-type (WT) or mutant tumors, that were generated as previously described^10^, Romidepsin treatment caused earlier and stronger suppression of growth in NRF2-active tumors (**Figures 6G-H, S6C)**, as well as reduction in c-MYC levels (**Figure S6D**). These results show that the phenotype and mechanism of HDAC inhibition we described in murine LUAD can be extrapolated to human settings and provide pre- clinical evidence of Romidepsin as a potential therapeutic for LUAD with NRF2-activation.

**Figure 6.**
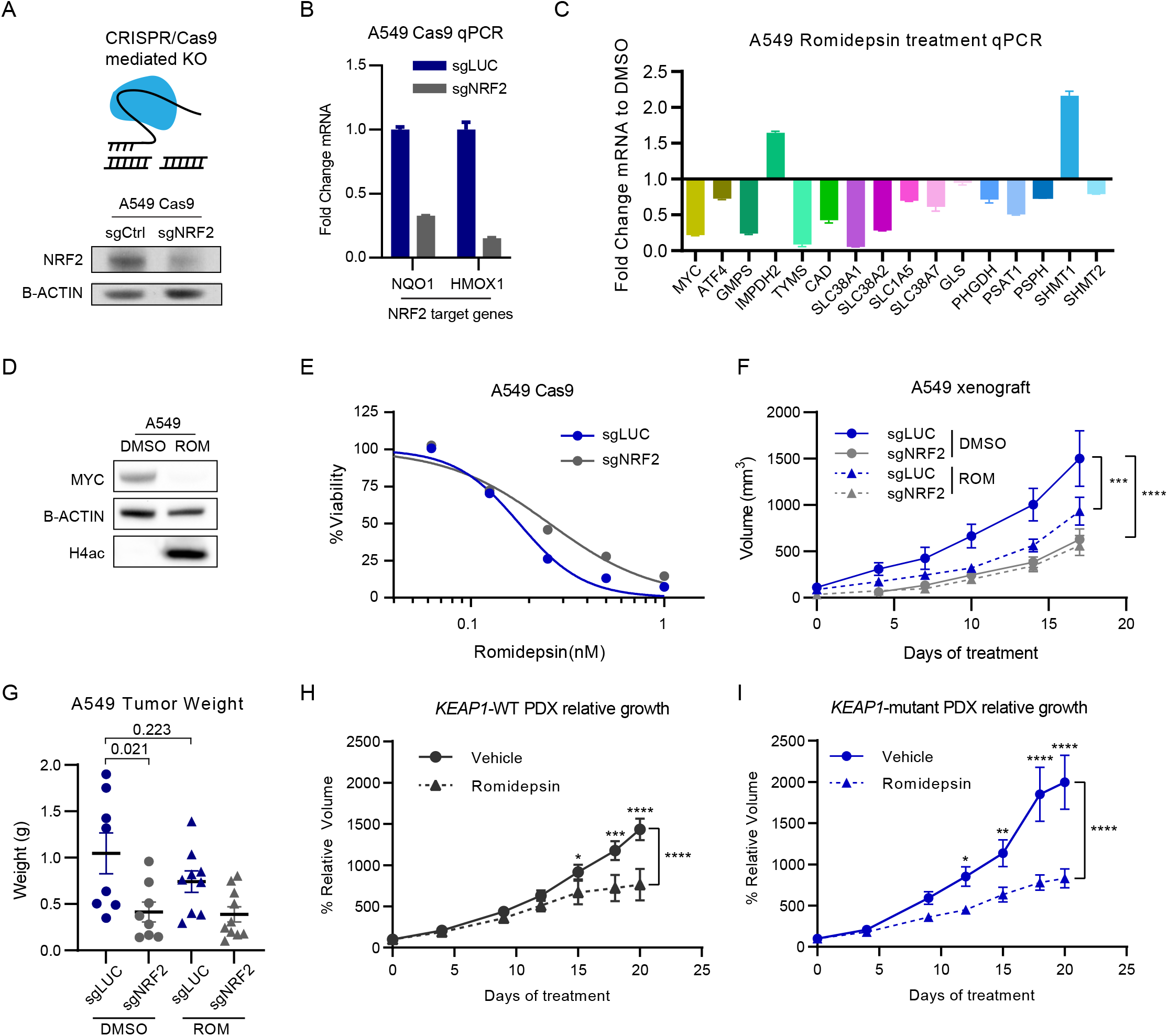
NRF2-activation confers sensitivity to HDAC inhibition in human cells and patient-derived xenografts (PDX). **A**. Schematic of CRISPR-mediated KO of NRF2, and NRF2 protein levels upon of NRF2 KO in a population of A549 cells (sgNRF2). **B**. Expression of NRF2 target genes upon NRF2 KO in A549 cells. **C**. Fold-Change expression of indicated genes upon Romidepsin treatment of A549 cells determined by RT-qPCR. **D**. Western blot showing levels of MYC and H4ac upon Romidepsin treatment of A549 cells 0.5nM for 24h. **E**. Viability assay of A549 cells treated with the indicated concentrations of Romidepsin for 72h. **F**. Growth of subcutaneous A549 tumors in nude mice treated with Romidepsin or vehicle, and tumor weight at the end of treatment (**G**). **H-I**. Relative growth curves of PDX tumors that are wild-type or mutant for *KEAP1* treated with vehicle or Romidepsin.

## Discussion

In this study, we used a focused CRISPR/Cas9 genetic screen to identify chromatin vulnerabilities driven by NRF2 activation in LUAD. We identified a preferential dependency on Class I HDAC genes, which translated to increased sensitivity to the Class I HDAC inhibitor Romidepsin. Following HDAC inhibition, global hyperacetylation of the genome displaces transcription co-activators such as BRD4 from genes with high levels of histone acetylation, such as those involved in cell metabolism. As a result, we observed reduced rate of glutamine uptake/catabolism as well as *de novo* nucleotide synthesis which imposed selective metabolic stress on NRF2-active cancer cells, causing anti-tumor effects that can be observed *in vitro* and *in vivo* using human NSCLC cell lines and patient-derived xenografts.

These preclinical findings have several mechanistic and translational implications. First, targeting of KEAP1/NRF2 alterations in cancer remains a key clinical priority. NRF2 hyperactivation promotes aggressive tumor growth and resistance to chemotherapy, radiation and immunotherapy, leading to poor patient prognosis. A number of therapeutic strategies have been explored, including small molecules targeting NRF2 itself or its interaction with KEAP1^32, 33^, tumor immune-modulating agents^34, 35^, and inhibiting metabolic pathways as synthetic lethal events^10, 11, 27, 36^. In particular, inhibitors of glycolysis and glutaminolysis have shown promising effects in preclinical models^10, 36, 37^, as NRF2 activation causes significant changes to central carbon and amino acid metabolism. Nevertheless, the glutaminase inhibitor CB-839 as monotherapy has shown limited success in treating *KEAP1* mutant NSCLC in clinical trials. Our study suggests that repurposing the FDA-approved HDAC inhibitor Romidepsin could represent another synthetic lethal approach to target NRF2-active tumors. Our *in vivo* study suggests that Romidepsin has similar efficacy as CB-839 and the combination of both inhibitors offers additional therapeutic effects. These results provide strong preclinical rationale for evaluating Romidepsin, alone or combined with CB-839, for treating NSCLC or other tumor types harboring dysregulated KEAP1/NRF2 pathway. In addition to glutamine metabolism, our epigenome and transcriptome analysis also identified *de novo* nucleotide synthesis as a novel metabolic vulnerability of NRF2-active cancer cells, which was confirmed by metabolomic and functional studies. Future efforts are required to evaluate the efficacy of inhibiting nucleotide synthesis, such as inhibitors of dihydroorotate dehydrogenase (DHODH), in treating NRF2 hyperactive tumors.

Mechanistically, our multi-omics analysis revealed that the impact of Romidepsin on histone acetylation, particularly H4ac, is more complex than previously reported. Specifically, the genome-wide modest gain in diffuse H4ac signal drives a relative loss of H4ac at strong promoter peaks. This redistribution in H4ac was associated with concomitant changes in BRD4 binding and gene expression. These findings are in agreement with a recent report^24^ and suggest that the initial chromatin state can be predictive of the epigenetic and transcriptional effects of HDAC inhibition. Transcriptomic and metabolomic analysis indicated that HDAC inhibition suppressed metabolic gene expression and activity in LUAD cells. Specifically, we found disruption of pathways that support viability and growth during NRF2 activation, including purine and pyrimidine synthesis, glutamine uptake and hydrolysis, serine synthesis, and the TCA cycle. The paradigm of metabolic gene expression modulation by epigenetic perturbation has been described in several other contexts, including *KMT2D*-deficient lung cancer^38^, H3K27- mutant glioma^39^ and childhood posterior fossa group A ependymomas^40^. In addition, dependency on transcriptional regulation of serine and nucleotide synthesis has been described in AML^41^. Future studies are warranted to investigate how chromatin abnormality can be hijacked by cancer cells to transcriptionally reprogram metabolism and if combined targeting of chromatin and metabolism represents an effective therapeutic strategy in additional settings.

Cancer-associated chromatin abnormality is emerging as a major target for therapeutic intervention, and specific and potent inhibitors of many chromatin-modifying enzymes have been developed^42^. Nevertheless, only a handful of chromatin-targeted drugs have been approved to treat mainly hematologic malignancies, including HDAC inhibitors (Romidepsin and Vorinostat) which are used for treatment of refractory T-cell lymphoma^43, 44^, as well as DNA methyltransferase inhibitors which are used as first-line treatment of myeloid malignancies^45^. In part, this gap reflects our limited understanding of the mechanisms of action and the lack of robust biomarkers for epigenetic therapies^46^. Previous studies assessed the efficacy of HDAC inhibitors mainly through phenotypic observations, gene expression analysis and analogue-based drug discovery^47^. As a result, their precise mechanism of cell killing is not well understood. Our findings indicate that Romidepsin transcriptionally reprograms amino acid metabolism and *de novo* nucleotide synthesis which renders selective toxicity to NRF2-active cancer cells. This is consistent with previous reports that HDAC inhibitors synergize with inhibitors of electron transport chain and fatty acid oxidation in glioblastoma^29^. Moreover, disruption of metabolic pathways by HDAC inhibitors has been reported by several other groups^48–50^. Taken together, these results suggest that metabolic alterations such as NRF2 hyperactivation could serve as effective biomarkers to predict the efficacy of HDAC inhibitors for treating solid tumors. We believe that similar concepts and strategies may be applicable to the identification of new biomarkers and broaden the number of cancer patients who could benefit from other epigenetic drugs.

In summary, our findings provide the evidence and mechanistic basis for Class I HDACs as potent and specific chromatin vulnerabilities of tumors with NRF2 activation. Moreover, they advocate retrospective or prospective studies on the NRF2 pathway as biomarkers to predict solid tumors’ response to HDAC inhibitors. We propose that the development of effective epigenetic therapies requires rational design of pre-clinical studies and patient stratification that consider the interplay between the genetic, chromatin and metabolic state of the cancer cell.

## Methods

### Cell culture

KP and KPK cell lines were established as described previously^10^. For experiments outlined in **Figures 1 and S1**, two independent KP and two independent KPK cell lines were used. For all other experiments, KP and KPK cells refer to one of the two KP and KPK cell lines respectively and n refers to the number of experimental replicates. All cells were maintained in either DMEM or RPMI-1640 (Sigma-Aldrich) supplemented with 10% FBS (Sigma-Aldrich) and 1x Penicillin-Streptomycin (Sigma-Aldrich). Cells were incubated at 37 degrees in 5% CO2 atmosphere. All cell lines were routinely tested for mycoplasma contamination. For antibiotic-based selection, puromycin (Sigma-Aldrich) was used at 5ug/ml, hygromycin (Sigma-Aldrich) at 500ug/ml and blasticidin (Sigma-Aldrich) at 5ug/ml. For fluorophore-based selection, cells were sorted by the Flow Cytometry Shared Resource at Columbia University using a BD Influx Cell Sorter.

### Plasmid construction and Lentivirus production

Lentivirus were generated by transfecting 293T cells with the indicated expression plasmid and the psPAX2 (Addgene) and pVSVG (Addgene) packaging vectors at a ratio of 4:2:3, respectively. Viral supernatants were collected 48 and 72 hrs after transfection, filtered and used for transduction of cells in 1:1 ratio with medium. NRF2ΔNeh2, Keap1 (mouse and human), and KEAP1^R470C^ overexpression constructs were generated by the Papagiannakopoulos lab. For CRISPR/Cas9 gene knock-out, we used the lentiCas9-blast plasmid (Addgene) and the pUSEPR vector for sgRNA (U6-sgRNA-EFS-Puro-P2A-TurboRFP in pLL3-based lentiviral backbone). For sgRNA design the CRISPick platform (BROAD institute) was used (**Supplementary Table 2**). For Slc1a5 overexpression, cDNA was obtained from Addgene (Plasmid #71458) and cloned into pLVX-IRES-mCherry (Takara Bio).

### sgRNA library

The gRNA library targeting murine chromatin regulators was constructed as previously described^51^. Briefly, it consisted of sgRNA sequences (six per gene) targeting 612 mouse chromatin regulators that were designed using BROAD sgRNA Designer^52^ and 36 non-targeting control sgRNAs^15^.This library was synthesized by Agilent Technologies and cloned into the pUSEPR lentiviral vector to ensure a library representation of >10,000X using a modified version of a previously described protocol^52^. Then, it was selectively amplified using barcoded forward and reverse primers that append cloning adapters at the 5 - and 3 -ends of the sgRNA insert, purified using the QIAquick PCR Purification Kit (Qiagen), and ligated into BsmBI-digested and dephosphorylated pUSEPR vector using high-concentration T4 DNA ligase (NEB). Ligated pUSEPR plasmid DNA was electroporated into Endura electrocompetent cells (Lucigen), recovered for one hour at 37⁰C, plated across four 15cm LB-Carbenicillin plates (Teknova), and incubated at 37⁰C for 16 hours. The total number of bacterial colonies was quantified to ensure a library representation of >10,000X. Bacterial colonies were scraped and briefly expanded for 4 hours at 37⁰C in 500mL of LB-Carbenicillin. Plasmid DNA was isolated using the Plasmid Plus Maxi Kit (Qiagen).

### Focused CRISPR/Cas9 Screen

Derivatives of KP and KPK cells were generated by stable lentiviral transduction of Cas9 with blasticidin resistance (Addgene#52962). Cells were maintained with blasticidin selection throughout the experiment. Transduction of Cas9-expressing KP/KPK cells was performed at an MOI of approximately 0.2 by incubating cell suspension in lentiviral supernatant and centrifugation at 1500 rpm for 1.5 hours at room temperature before being returned to a humidified incubator. An initial population was infected to represent a 2500x representation of the epigenetic library; 36 hours post-transduction cells were resuspended and replated in 10 ug/mL puromycin and selected for another 48 hours. After complete puromycin selection cells were trypsinized, pooled, a cell sample representing time-0hr (t_0_) of the screen was reserved and stored at −20⁰C. Each condition was performed in technical triplicate for the entire screen and maintained in 15 cm tissue culture dishes (Corning) with at least 2500x library representation maintained throughout all culture and library preparation steps. Population doublings for each cell line and condition were recorded and a sample was collected when a particular condition had reached 14 cumulative population doublings and stored at −20⁰C.

Genomic DNA from collected cell pellets were prepared with purelink mini kit (ThermoFisher) according to the manufacturer’s suggested protocol. Amplification of 5ug gDNA equivalent was done as described previously^51, 53^ and sequenced using an Illumina Nextseq 500 high output with 40% PhiX spike-in.

Computational analysis was done using MAGeCK^54^. Briefly, the sequencing data were de-barcoded and the 20 bp sgRNA sequence was mapped to the reference sgRNA library without allowing for any mismatches. The read counts were calculated for each individual sgRNA and normalized, and differential analysis was done between KP and KPK samples. Quality control, gene hit identification and graphs were generated using MAGeCKFlute^55^.

### Cell growth competition assay

Cells were transduced with Cas9, selected with blasticidin for 1 week, and then transduced with the sgRNA constructs containing RFP overexpression (at least 2 per gene). At 3 days (t_0_) and then after 14 population doublings (t_1_), cells were analyzed by flow cytometry. Flow cytometry data was acquired on 16 laser BDFortessa. All data were analyzed using the FlowJo^TM^ (V10) software. The percentage of RFP positive cells was determined by gating using uninfected Cas9 cells for each cell line.

### Protein Extraction and Western blot analysis

Whole cell lysates were prepared in SDS Lysis Buffer (ThermoFisher) and resolved on 3-8% or 4-12% gradient SDS-PAGE gels (ThermoFisher) transferred to nitrocellulose membrane, blocked in 5% non-fat milk in PBS plus 0.5% Tween-20, probed with primary antibodies (**Supplementary Table 3**) and detected with horseradish peroxidase-conjugated α-rabbit or α-mouse secondary antibodies (Cell Signaling). The blots were imaged using a ChemiDoc MP Imaging system (Bio-Rad) or exposed to X-Ray films (Research Products International).

### Viability assays and drug treatments

For 72h viability assays, 1500 KP cells, 3000 A549 or 5000 H2009 cells were plated in 96-well plates in RPMI-1640 medium, the next day cells were treated (**Supplementary Table 4**), and cell viability was determined 72h post treatment using CCK8 assay (Dojindo). AUC and IC50s were determined using the Graphpad Prism v6 and v9 software. For 5-day assays, 1500 KP or KPK cells were plated in 12-well plates in RPMI-1640 medium, the next day cells were treated (**Supplementary Table 4**), and cell viability was determined post treatment by Crystal Violet stain (Sigma). For induction of NRF2 activation in KP carrying dox-inducible NRF2ΔNeh2, cells were treated with 1µM KI-696 (Papagiannakopoulos lab) or 1µg/ml Doxycycline (Sigma-Aldrich) for 7 days before experiments.

The concentration of *in vitro* Romidepsin treatments throughout the study varies due to use of two stocks with different potency (second batch of drug was 5 to 10-fold more potent), as well as to account for differences in cell number and plate well size. For specific experiments the concentrations used were: CUT&Tag and RNA-seq: 5nM for 17h, ^13^C-glucose tracing experiment: 5nM, ^13^C-glutamine tracing experiment: 1nM as indicated in **Figure S5A**.

### Allograft and Xenograft studies

For KP and A549 *in vivo* studies, 6-8 weeks old male C57BL/6 mice (Cat# 000664) and *Foxn1^nu^* mice (Cat# 007850) were purchased from The Jackson Laboratory. All mice were housed under specific-pathogen-free (SPF) conditions and followed the guidelines of Columbia University animal facility. All mice experiments were carried out with the protocol approved by the Institutional Animal Care and Use Committee (IACUC) at Columbia University. C57BL/6 mice or *Foxn1^nu^* mice were subcutaneously injected with KP (5 × 10^5^ per injection) and A549 (5 × 10^6^ per injection) cells respectively into the flanks (2 injections/mouse). Mice were treated after tumor establishment, approximately 5 days post injection. Subsequent intraperitoneal (IP) treatments and tumor measurements were performed 2-3 times a week on the days indicated in each figure. Romidepsin (1mg/kg IP; Medchem Express) was dissolved in 10% DMSO in Corn oil (Sigma Aldrich). CB-839 (200mg/kg Orally; twice a day; CALITHERA) was formulated in 25% (w/v) hydroxypropyl-β-cyclodextrin in 10 mmol/L citrate (pH 2.0), at 20 mg/mL for a final dosing volume of 10 mL/kg.

For the Patient-derived xenograft (PDX) experiment, the study was approved by the NYU Langone Medical Center Institutional Animal Care and Use Committee. Animals were housed according to IACUC guidelines in ventilated cages in a specific pathogen-free (SPF) animal facility. PDX tumors were stored in cryo-tubes in 10% dimethyl sulphoxide (DMSO) containing Dulbecco’s Modified Eagle Medium (DMEM) media containing 10% FBS and 20 ug/ml Gentamicin. After stabilized and expanding in NOD-scid IL2R gamma null (NSG) mice, tumors were trimmed with the size of 3 mm x 3 mm x 3 mm and subcutaneously transplanted near both flanks into NSG male and female littermates approximately 6-8 weeks in age. Engraftment was checked every 5 days after transplantation. After the tumor establishment phase, animals were randomized and assigned to a treatment group. Tumor volume was measured by caliper and volume was calculated (Length x Width^2^ x 0.5). Animals either received Romidepsin 1 mg/kg or vehicle Corn Oil twice weekly administered through intraperitoneal injection. The treatment volume was settled as 100 µl per mouse. Tumor growth was tracked for a minimum of 8 tumors per experimental group. Tumors with volume less than 20mm^3^ at the time of the first measurement were excluded from the final analysis.

Statistical analyses were done using Prism (v9), specifically 2-way ANOVA was used for comparison of tumor growth between each condition and Fisher’s Least Significant Difference for multiple comparisons. For comparisons of tumor volumes and weights, the test was chosen by performing D’Agostino-Pearson and Shapiro-Wilk normality test: if both conditions passed both tests, Student’s t-test was used, otherwise we performed Mann-Whitney U-test.

### CUT&Tag

CUT&Tag was performed as described previously^23^, with an additional step of light fixation to better preserve histone acetylation/TF binding. In brief, 1 × 10^5^ cells were lightly fixed with 0.1% paraformaldehyde 5’, neutralized by Glycine 125mM, and washed once with 1 ml of wash buffer (20 mM HEPES pH 7.5, 150 mM NaCl, 0.5 mM Spermidine (Sigma-Aldrich), 1× Protease inhibitor cocktail (Roche)). Concanavalin A-coated magnetic beads (Bangs Laboratories) were washed twice with binding buffer (20 mM HEPES pH 7.5, 10 mM KCl, 1 mM MnCl2, 1 mM CaCl2). 10 µl/sample of beads were added to cells in 400ul of wash buffer and incubated at room temperature for 15 min. Beads-bound cells were resuspended in 100 µl of antibody buffer (20 mM HEPES pH 7.5, 150 mM NaCl, 0.5 mM Spermidine, 0.06% Digitonin (Sigma-Aldrich), 2 mM EDTA, 0.1% BSA, 1× Protease inhibitor cocktail and incubated with indicated antibodies (**Supplementary Table 3**) or normal rabbit IgG (Cell Signaling) at 4 degrees overnight on nutator. After being washed once with Dig-wash buffer (20 mM HEPES pH 7.5, 150 mM NaCl, 0.5 mM Spermidine, 0.05% Digitonin, 1× Protease inhibitor cocktail), beads-bound cells were incubated with 1 µl Guinea pig anti-rabbit secondary antibody (Antibodies Online ABIN101961) and 2 µl Hyperactive pA-Tn5 Transposase adapter complex in 100 µl Dig-300 buffer (20 mM HEPES•NaOH, pH 7.5, 0.5 mM Spermidine, 1× Protease inhibitor cocktail, 300 mM NaCl, 0.01% Digitonin) at room temperature for 1 h. Cells were washed three times with Dig-300 buffer to remove unbound antibody and Tn5 and then resuspended in 300 µl of tagmentation buffer (10 mM MgCl2 in Dig-300 buffer) and incubated at 37 °C for 1 h. 10 µl of 0.5 M EDTA, 3 µl of 10% SDS and 5 µl of 10 mg/ml Proteinase K were added to each sample and incubated at 50 °C for 1 h to terminate tagmentation. DNA was purified using chloroform isoamyl alcohol (Sigma Aldrich) and eluted with 25 µl ddH2O. For library amplification, 21 µl of DNA was mixed with 2µL i5 unique index primer (10 µM), 2 µL i7 unique index primer (10 µM) and 25 µL NEBNext High-Fidelity 2X PCR Master Mix (NEB) and subjected to the following PCR program: 72℃, 5 min; 98℃, 30 sec; 13 cycles of 98℃, 10 sec and 63℃, 10 sec; 72℃, 1 min and hold at 10℃. To purify the PCR products, 1.1× volumes of pre-warmed Ampure XP beads (Beckman Coulter) were added and incubated at room temperature for 10 min. Libraries were washed twice with 80% ethanol and eluted in 20 µl of 10 mM Tris-HCl, pH 8. Libraries were sequenced on an NextSeq 550 platform (Illumina, 75 cycles High Output Kit v2.0) and 75-bp paired-end reads were generated. To determine global level differences in histone acetylation signal we used spike-in controls. For H3K27ac, 2ul of SNAP-ChIP K-AcylStat panel nucleosomes (EpiCypher) was added as spike-in control at the primary antibody incubation step. For H4ac, 5000 S2 Drosophila cells were added at the cell-bead binding step.

### CUT&Tag data analysis

CUT&Tag reads of KP cell samples were mapped to the mouse genome assembly mm10 using Bowtie2 (v2.3.5.1, parameters: --local --very-sensitive-local --no-unal --no-mixed --no-discordant --phred33 -I 10 -X 700). Potential PCR duplicates were removed by the function “MarkDuplicates” (parameter: REMOVE_DUPLICATES=true) of Picard (v2.24.2). Genomic enrichments of CUT&Tag signals were generated using deeptools (v3.3.2, parameters bamCoverage --normalizeUsing CPM --binSize 25 --smoothLength 100) and visualized using IGV. Peaks were called using MACS2 (parameters: --f BAMPE -g mm --broad). Consensuses of H3K27ac, H4ac and BRD4 peaks across conditions were generated by the ‘cat’ function (Linux) and ‘sort’ and ‘merge’ functions of bedtools (v2.27.1). The read counts of H3K27ac, H4ac and BRD4 CUT&Tag data in genomic elements were measured by featureCounts (v2.0.0). Differential analysis was performed using DEseq2(v1.32.0). Peak annotation was done using ChIPseeker^56^(v1.28.3). Heatmaps were generated using deeptools (v3.3.2) functions computeMatrix and plotHeatmap. For visualization we used the R package ggplot2 (v3.3.2). For genome-wide signal difference correlations we used deeptools (v3.3.2) functions multiBigwigSummary and plotCorrelation. Promoters were defined as 2.5kb regions centered around the TSS. For the signal diffusion analysis, random regions were generated using the bedtools(v2.27.1) ‘shuffle’ function and the called peaks as input. For the assignment of genes into quantiles (Figures 3E and 3G), genes were ranked by their promoter CPM and split into percentiles (3E: 0-33, 33-66 and 66-100, 3G as labeled).

To determine global level differences in histone acetylation we compared the ratio of spike-in reads to total number of reads. For K27ac, the number of reads for each barcode was counted to determine the scaling factor. For H4ac, reads were mapped to the Drosophila genome (Dmel_A4_1.0) and the mapping percentage was used to determine the scaling factor.

### RNA isolation, quantitative reverse transcription PCR (RT-qPCR) and RNA-sequencing

Total RNA was extracted in TRIzol (Invitrogen) and precipitated in ethanol (DECON Labs). For qRT-PCR, cDNA was then synthesized with cDNA Synthesis Kit (Takara) according to the manufacturer’s protocols. The relative expression of targeted genes was measured by qRT-PCR with indicated primers and SYBR Green Master Mix (ThermoFisher) using the ABI 7500 Real-Time PCR Detection System (Applied Biosystems). The sequences of primers used are listed in (**Supplementary Table 5**). For RNA-sequencing, RNA samples were submitted to Columbia University Genome Center for library preparation, sequencing and bioinformatic analysis up to generation of a reads count table of each gene. The differential gene expression was calculated by the R package DESeq2 (v1.28.0), and visualization was done using ggplot2 (v3.3.2),Pretty Heatmaps (v1.0.12) and Enhanced Volcano (1.10) R packages.

### DepMap dataset analysis

The datasets that were used were the CERES 21Q3 Public+Score and the Prism repurposing secondary screen 19Q4. Data was downloaded for subsets that included NSCLC cell lines with or without KEAP1 mutations. For statistical analysis, the test was chosen by performing D’Agostino-Pearson and Shapiro-Wilk normality test: if both conditions passed both tests, Student’s t-test was used, otherwise we performed Mann-Whitney U-test. For CERES, comparison of the mutant and wild-type cell line subsets was done by calculating the gene-effect difference between the two.

### Metabolic tracing

For glucose tracing analysis, 2×10^5^ KP cells (n=3) were plated in 6-well plates overnight in RPMI medium (Sigma). After 24 hours, the media was replaced with fresh RPMI medium containing DMSO or 5nM Romidepsin. At 48 hours the media was replaced with fresh glucose-free RPMI medium (Sigma) containing 10% dialyzed fetal bovine serum (Gibco), 2.0 g/L ^13^C_6_- glucose (Sigma) and DMSO or 5nM Romidepsin. Cells were harvested at 49 and 72 hours and processed as described below.

For glutamine tracing, 2×10^5^ KP cells (n=3) were plated in 6-well plates overnight in RPMI medium (Sigma). After 24 hours, the media was replaced with fresh RPMI medium containing DMSO, 1nM Romidepsin or 150nM CB-839. At 48 hours the media was replaced with fresh glutamine-free RPMI medium (Sigma) containing 10% dialyzed fetal bovine serum (Gibco), 2.0 g/L ^13^C_6_-glutamine (Cambridge Isotope Laboratories) and DMSO, 1nM Romidepsin or 150nM CB-839. Cells were harvested at 56 hours and processed as described below.

### Metabolite harvesting and liquid chromatography-mass spectrometry analysis

Cells were washed with cold PBS, lysed in 80% Ultra LC-MS acetonitrile (Thermo Scientific) supplemented with 20 µM deuterated 2-hydroxyglutarate (D-2-hydroxyglutaric-2,3,3,4,4-d5 acid (d5-2HG), Cambridge Isotope Laboratories) as an internal standard on ice for 15 minutes, and centrifuged for 10 minutes at 20,000 x g at 4 °C. 200 µL of supernatants were subjected to mass spectrometry analysis. Liquid chromatography was performed using an Agilent 1290 Infinity LC system (Agilent, Santa Clara, US) coupled to a Q-TOF 6545 mass spectrometer (Agilent, Santa Clara, US). A hydrophilic interaction chromatography method with a ZIC-pHILIC column (150 x 2.1 mm, 5 µm; EMD Millipore) was used for compound separation at 35 °C with a flow rate of 0.3 mL/min. Mobile phase A consisted of 25 mM ammonium carbonate in water and mobile phase B was acetonitrile. The gradient elution was 0—1.5 min, 80% B; 1.5—7 min, 80% B → 50% B, 7—8.5 min, 50% B; 8.5—8.7 min, 50% B → 80% B, 8.7-13 min, 80% B. The overall runtime was 13 minutes, and the injection volume was 5 µL. The Agilent Q-TOF was operated in negative mode and the relevant parameters were as listed: ion spray voltage, 3500 V; nozzle voltage, 1000 V; fragmentor voltage, 125 V; drying gas flow, 11 L/min; capillary temperature, 325 °C; drying gas temperature, 350 °C; and nebulizer pressure, 40 psi. A full scan range was set at 50 to 1600 (m/z). The reference masses were 119.0363 and 980.0164. The acquisition rate was 2 spectra/s. Targeted analysis, isotopologues extraction (for the metabolic tracing study), and natural isotope abundance correction were performed by the Agilent Profinder B.10.00 Software (Agilent Technologies).

### Histology

Tumors were fixed in 10% Formalin (Fisher Chemical) for 48 hours and then stored in 70% ethanol at 4°C. Paraffin embedding and sectioning was done at the Histology Service of the Molecular Pathology Shared Resource at Columbia University Medical center. Immunohistochemistry experiments were done at Experimental Pathology Research Laboratory at New York University Langone Health. Quantitation of signal was done using QuPath software (v0.3.2) and statistical analysis using Prism (v9) and Fisher’s Least Significant Difference for multiple comparisons; at least 4 tumors per condition were assessed.

### Glutamine and glutamine consumption

2×10^5^ KP cells (n=3) were plated in 6-well plates overnight in RPMI medium (Sigma). After 24 hours, the media was replaced with fresh RPMI medium containing DMSO, 1nM Romidepsin or 150nM CB-839. At 48 hours the media was replaced with fresh glutamine-free RPMI medium (Sigma) containing DMSO, 1nM Romidepsin or 150nM CB-839. Media were harvested at 56 hours, centrifuged to remove dead cells and frozen at −80°C. Cells were harvested and counted. Measurement of metabolites was done using a YSI 7000 enzymatic analyzer at the Cell Metabolism core that is part of the Donald B. and Catherine C. Marron Cancer Metabolism Center at the Memorial Sloan Kettering Cancer Center. Consumption was calculated by comparison to media from wells without any cells.

## Author Contributions

D.K., T.P. and C.L. conceived the study. D.K. executed the experiments with the help of W.W (CRISPR/Cas9 focused genetic screen), A.L. (metabolomics), M.H. (PDX), X.C., X.X., M.Y and V.M. F.J.S.-R. and Y.M.S.F. provided reagents and guidance for the CRISPR/Cas9 genetic screen. T.P. and J.Y. provided reagents, expertise, and feedback. C.L. supervised the study.

D.K. and C.L. wrote the manuscript, with contributions and input from all authors.

## Acknowledgements

We thank members of the Lu lab for critical reading of the manuscript. We thank Tahir Sheikh and the Gary Swartz lab for sharing reagents. We thank Zhiming Li and the Zhiguo Zhang lab for assistance with library sequencing. We thank Jozef Piotr Bossowski for assistance with ATF4 expertise and IHC staining. This study was funded by NIH (R35GM138181 and R01DE031873 to C.L.). D.K. acknowledges support from NYSTEM training grant. Research reported in this publication was performed in the CCTI Flow Cytometry Core, supported in part by the Office of the Director, National Institutes of Health under awards S10OD020056. The content is solely the responsibility of the authors and does not necessarily represent the official views of the National Institutes of Health. Immunohistochemistry experiments performed by the NYU Experimental Pathology Research Laboratory were funded in part by the NYUCI Center Support Grant, “NIH/NCI 5 P30CA16087”. F.J.S.-R. was supported by the MSKCC TROT program (5T32CA160001), a GMTEC Postdoctoral Researcher Innovation Grant, and is an HHMI Hanna Gray Fellow. Y.M.S.F was supported by the Damon Runyon-Sohn Pediatric Cancer Fellowship (DRSG-21-17) and NIGMS-MOSAIC K99/R00 Career Development Award (1K99GM140265). J.Y. was supported by an American Cancer Society Research Scholar Grant (RSG-20-036-01).

## Declaration of interests

Authors declare no competing interests.

---

**Figure S1.**
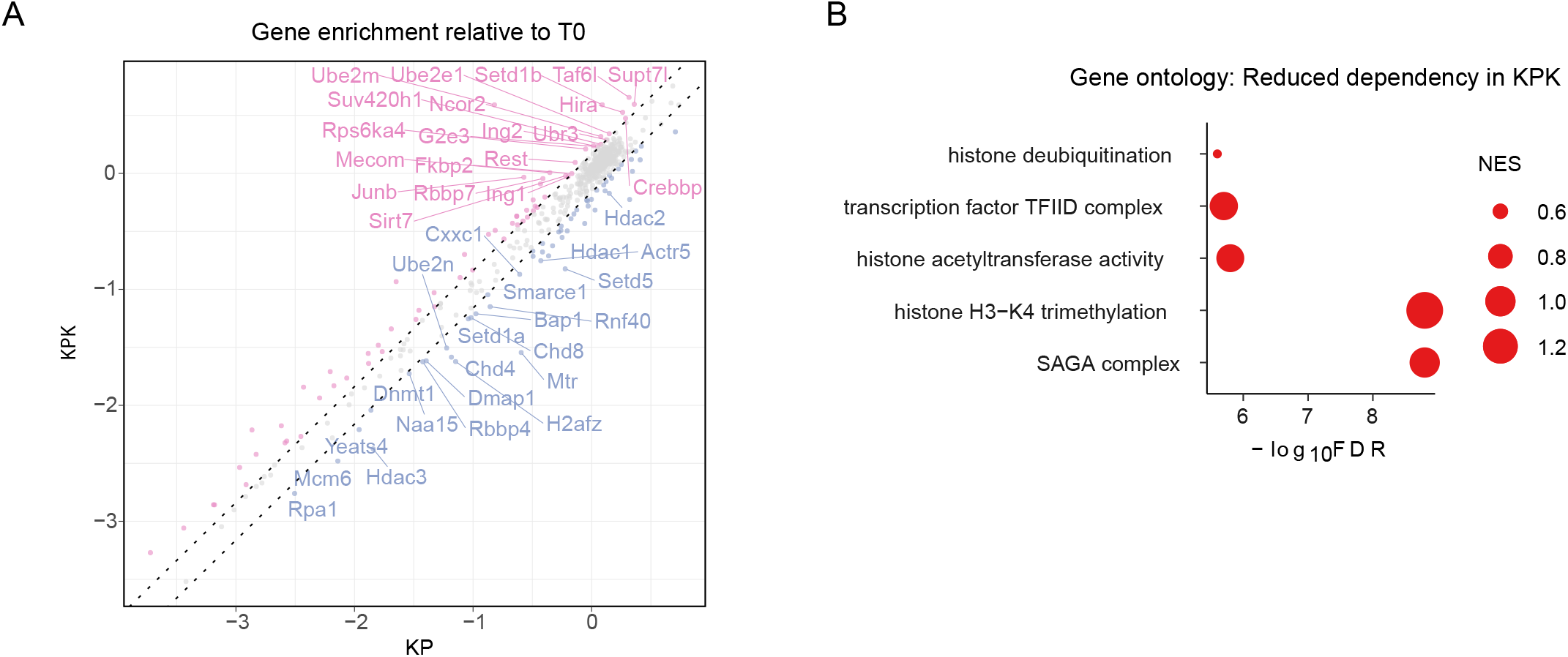
**A.** Scatter plot of gene enrichment compared to T0 for each condition and indicated are the top 20 differentially enriched genes. **B**. Ontology analysis of genes that are more essential in KP cells.

**Figure S2.**
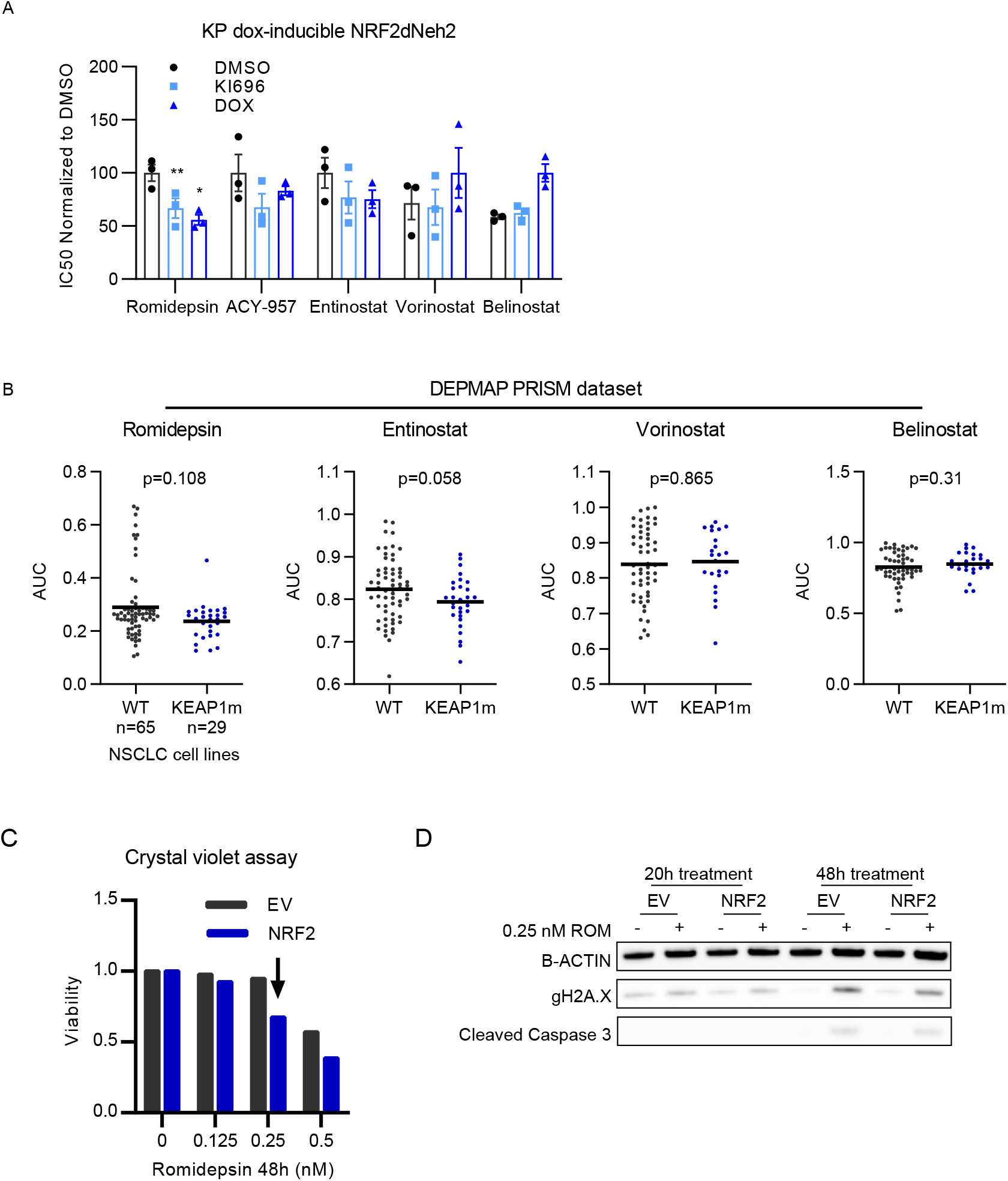
**A.** Bar graph indicating IC50s to various HDAC inhibitors, derived from 3-day viability experiments (n=3) in KP cells carrying dox-inducible NRF2ΔNeh2 pre-treated with DMSO, KI696 or DOX for 1 week (Statistical significance determined by paired t-test). **B**. HDAC inhibitor AUC (area under curve) comparison of KEAP1 WT and mutant NSCLC cell lines from the DepMap Prism repurposing secondary screen 19Q4. **C** and **D**. Protein levels of phosphorylated H2A.X and cleaved caspase 3 upon treatment with Romidepsin at a concentration were the difference in cell viability between control and NRF2 activated cells is more pronounced.

**Figure S3.**
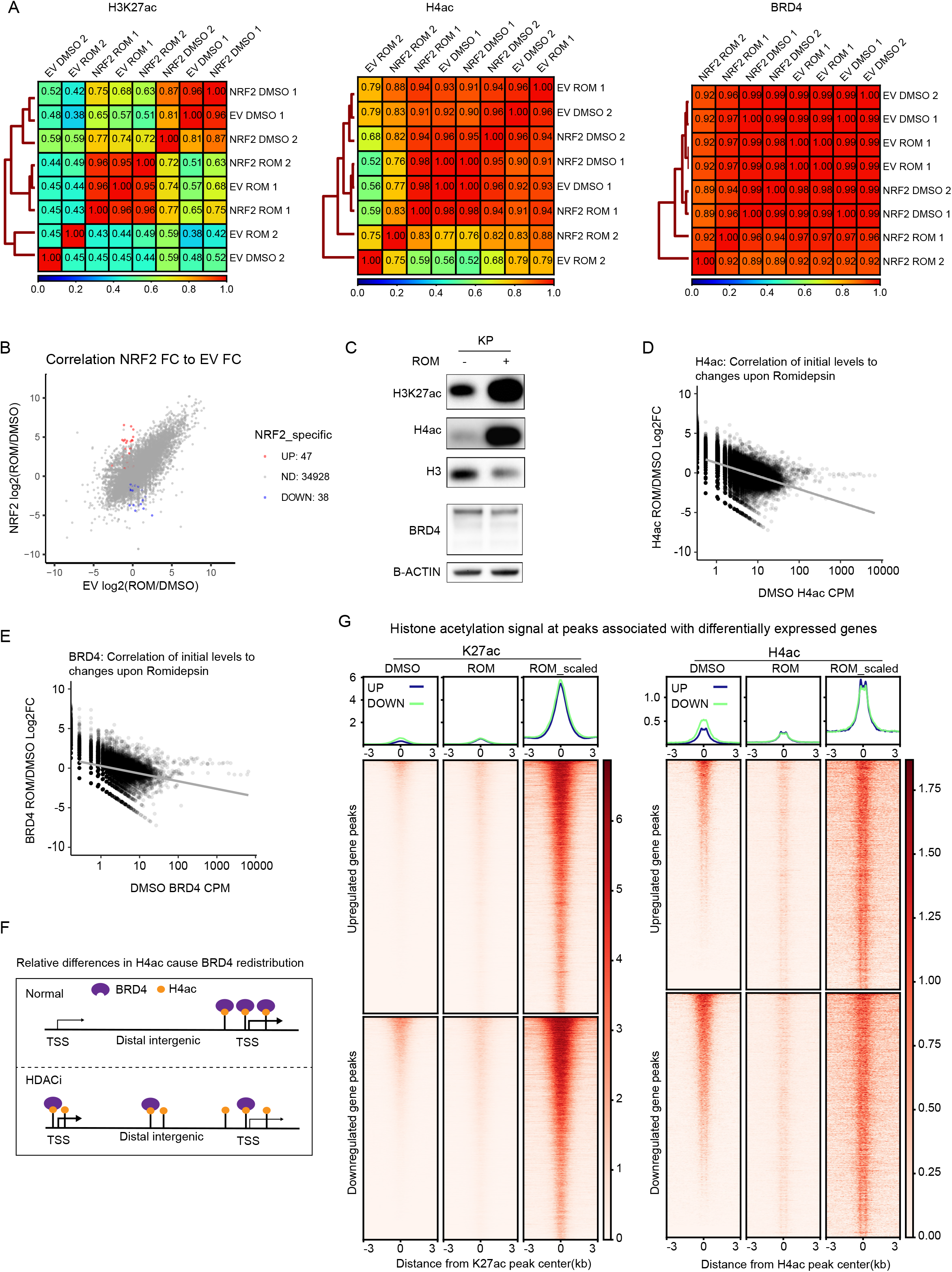
**A.** Pearson correlation heatmaps of all conditions and replicates. **B**, Correlation plot of Fold change differences between KP EV and NRF2 cells upon Romidepsin treatement; indicated are transcripts that are significantly up- or down-regulated only in NRF2 cells (cutoff: 2-fold change and FDR<0.05). **C.** Western blot analysis of H3K27 and H4 acetylation upon treatment of KP cells with Romidepsin (5nM, 24h). **D**. Correlation plot of fold-change histone acetylation and BRD4 binding (**E**) to levels of acetylation/binding at DMSO treatment at consensus peaks (normalized by DEseq2; n=2). **F**. Illustration showing how HDAC inhibition can cause gene expression changes by displacing and diffusion of BRD4 binding due to global gain of H4ac. **G**. Heatmaps of histone acetylation signal at peaks associated with differentially expressed genes in KP cells treated with DMSO or Romidepsin (scaled by RPKM or spike-in control).

**Figure S4.**
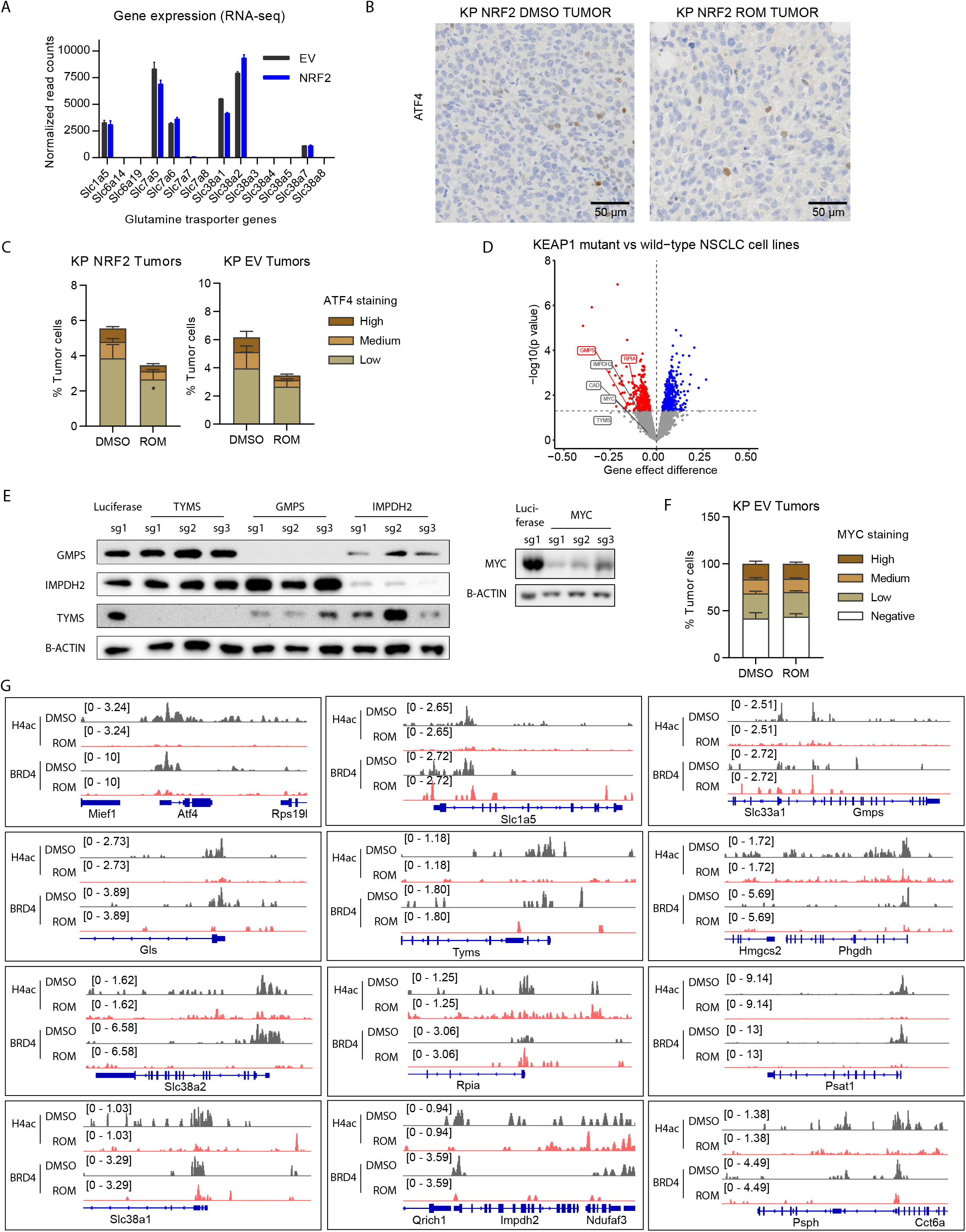
**A.** Normalized RNA-seq read counts (by DESeq2) of amino-acid transporters involved in glutamine uptake. **B**. IHC staining of ATF4 and quantitation (**C**) in KP tumors after 17 days of treatment with DMSO or Romidepsin (related to figure 2G-H). **D**. Volcano plot showing gene effect difference between KEAP1 wild-type and mutant NSCLC cell line from the CERES DepMap dataset. **E**. Western blots indicating levels of indicated proteins upon CRISPR/Cas9 KO in KP cells - associated with competition assay in 4F. **F**. Quantitation of IHC staining of c-MYC in KP EV tumors (related to figure 2G-H). **G**. Bedgraphs of H4ac and BRD4 binding at indicated genes.

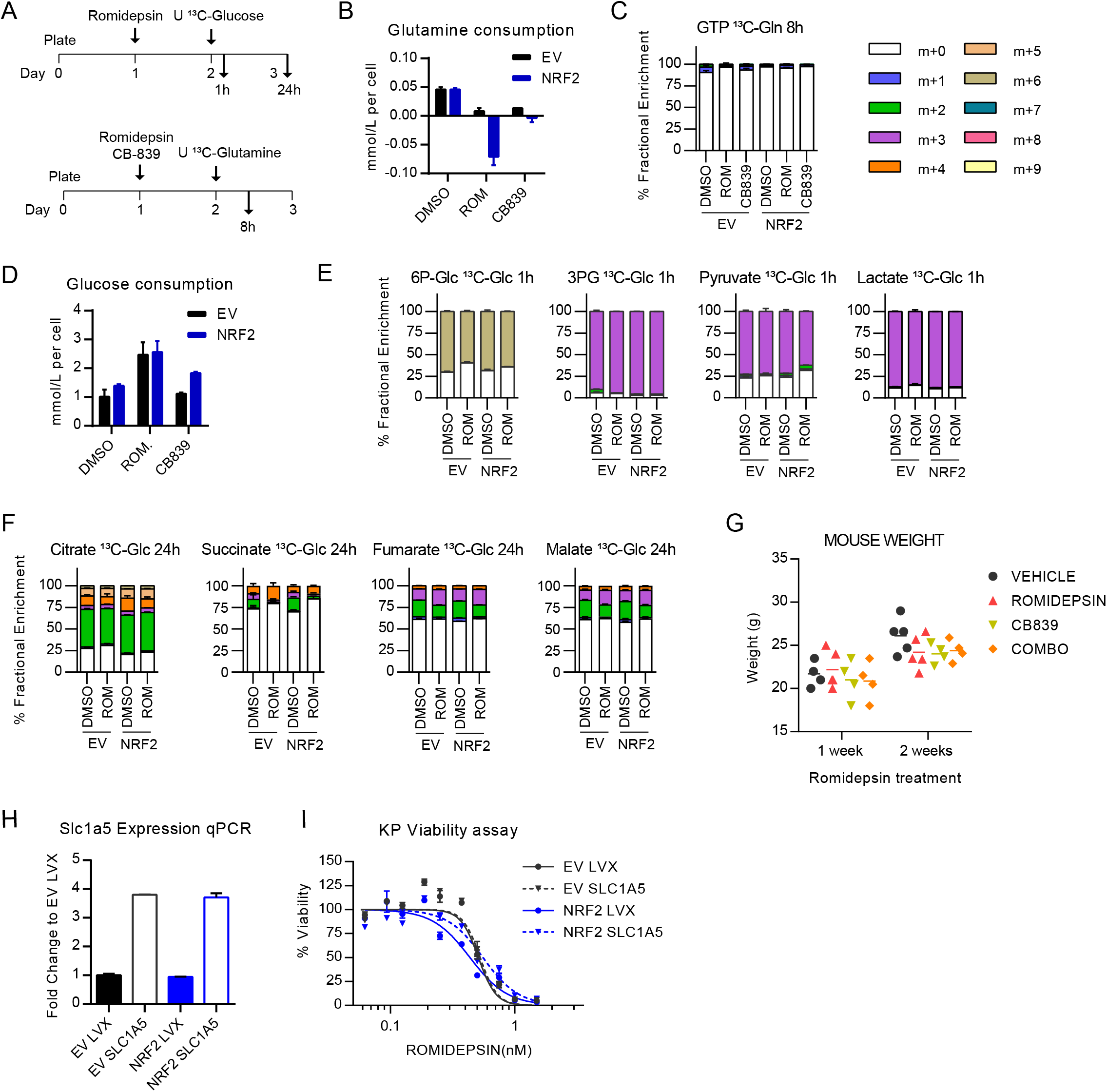

**Figure S6.**
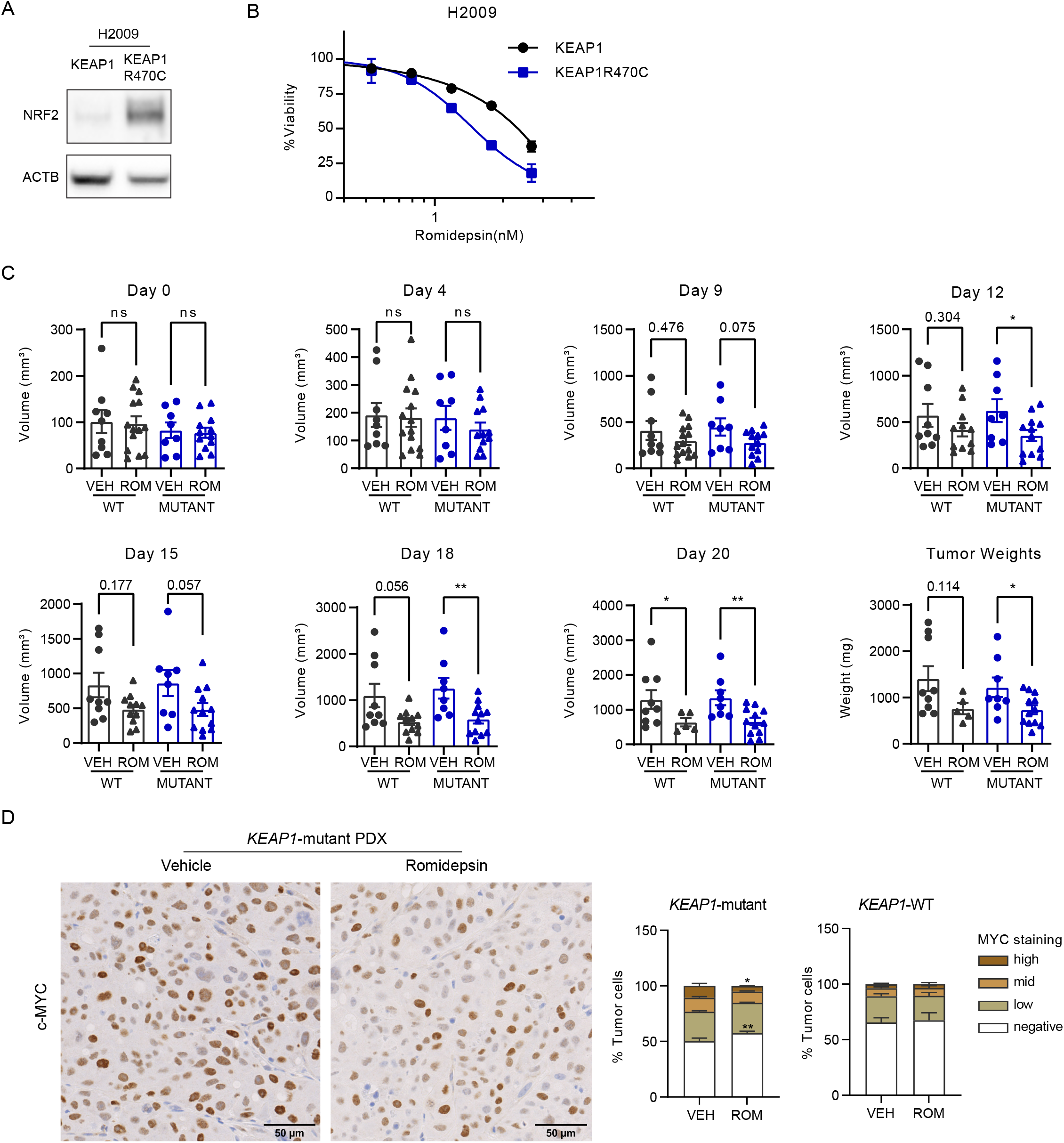
**A.** NRF2 protein levels upon overexpression of wild-type or R470C mutant KEAP1 in NCI-H2009 cells. **B**. Viability assay of H2009 cells treated with the indicated concentrations of Romidepsin for 72h. **C**. Tumor volumes of PDX tumors at the indicated days of treatment and tumor weights at the experiment endpoint (related to 6H-I). **D**. IHC staining of c-MYC and quantitation in PDX tumors after 17 days of treatment with Vehicle (VEH) or Romidepsin (related to figure 6H-I).

## Notes

### Competing Interest Statement

The authors have declared no competing interest.

